# Exploring Mechanisms of Inhibition of Amyloid Seeding of Transthyretin

**DOI:** 10.1101/354720

**Authors:** Lorena Saelices, Kevin Chung, Ji H. Lee, Teresa Coelho, Johan Bijzet, Merrill D. Benson, David S. Eisenberg

## Abstract

Amyloid deposition of the hormone transporter transthyretin causes familial and sporadic amyloidoses. The current treatment for familial cases is gene-therapy by liver transplantation. However, this procedure is often insufficient to stop subsequent cardiac deposition. Our recent work has shown that preformed amyloid fibrils present in the heart by the time of surgery can template or *seed* further polymerization of native transthyretin. No drugs have been approved to stop or slow this seeding process; the only treatment option is heart transplantation. Here we explore two potential inhibitory mechanisms. Of clinical significance, we found that tetramer stabilization does not hinder amyloid seeding. In contrast, binding of the peptide inhibitor TabFH2 to *ex-vivo* fibrils efficiently inhibits amyloid seeding in a tissue-independent manner. Our findings point to inhibition of amyloid seeding by peptide inhibitors as a potential therapeutic approach to be further explored.

## Introduction

Transthyretin amyloidosis (ATTR) is a fatal disease caused by the abnormal aggregation of the protein transthyretin (TTR). TTR is mainly secreted by the liver and choroid plexus, and transports retinol and thyroxine in blood and cerebrospinal fluid. TTR amyloid deposits are found in virtually every organ of ATTR patients, although the heart and nerves are often the first to fail. Although more than 140 known familial mutations in the *ttr* gene result in an early onset of the disease, wild-type TTR is found not only co-depositing with mutant TTR in hereditary ATTR cases but also in sporadic cases, in which the only TTR variant is wild-type. Sporadic ATTR, also known as wild-type ATTR or senile systemic amyloidosis, manifests as an aging-related disease, and is often overlooked and underdiagnosed (1, 2).

The current standard of care for hereditary cases worldwide is liver transplantation. Through this procedure, most of circulating mutant TTR is replaced with the wild-type form that is secreted by the implanted liver. However, this extreme surgery is not sufficient to stop amyloid cardiac deposition in many patients that end up requiring secondary heart transplantation a few years later. Our recent studies have shown that pre-formed TTR fibrils present in the heart of ATTR patients by the time of surgery have the capacity to catalyze or *seed* fibril polymerization of wild-type TTR that is secreted by the implanted liver (3).

The stabilization of the functional non-amyloidogenic form of transthyretin is currently under clinical assessment. The functional and most abundant form of TTR is tetrameric, with a hydrophobic central tunnel that binds thyroxine. Kelly and colleagues have established that conversion of native transthyretin to amyloid fibrils is preceded by dissociation of tetrameric TTR to monomers, which then undergo a conformational change and form fibrils (4). Based on this premise, extensive biochemical studies have led to the discovery of compounds such as tafamidis and diflunisal that bind within the hydrophobic central tunnel of TTR and stabilize the native structure, inhibiting its aggregation *in vitro* (5–7). These two ligands stabilize tetrameric transthyretin *in vivo* and delay progression of disease in many patients. However, the efficacy of such ligands is reduced when administered at late stages of the disease (8, 9).

In our recent studies, we have developed and optimized peptide inhibitors that are designed to cap the tip of TTR fibrils and block further polymerization (3, 10). This structure-based drug design strategy started with the identification of two amyloid-driving segments of transthyretin that comprise the β-strands F and H (10). We then determined the structures of the two segments in their amyloid state and designed peptide inhibitors that block self-association and protein aggregation *in vitro*. This first generation of inhibitors was further optimized in a second study (3).

Amyloid seeding represents a potential therapeutic target to be explored. Clinical observations indicate that amyloid seeding greatly contributes to ATTR pathogenesis. For instance, inhibiting amyloid seeding may potentially hinder post-surgical ATTR cardiac deposition after liver transplantation. In our previous study, we found that tetramer stabilizers do not inhibit amyloid seeding catalyzed by amyloid fibrils extracted from an ATTR-D38A patient (3). In contrast, peptide inhibitors halted this process. Here we expand our studies and evaluate the efficacy of tetramer stabilization and peptide inhibitors in blocking amyloid seeding of ATTR fibrils extracted from several patients and tissue types.

## Results

### *Tetramer stabilization by ligands does not inhibit TTR amyloid formation induced by ATTR-D38* exvivo *seeds*

We previously found that the presence of tafamidis or diflunisal at 180 µM is not sufficient to prevent amyloid seeding of wild-type TTR catalyzed by ATTR fibrils extracted from an ATTR-D38A patient (3). Here we expand the study by evaluating the effect of these ligands at various concentrations.

We first confirmed the ability of tafamidis and diflunisal to inhibit protein aggregation in the absence of seeds. Consistent with previous studies, we found that both tafamidis and diflunisal efficiently inhibit wild-type TTR aggregation *in vitro* under acidic conditions in the absence of seeds (Fig. 1a,b) (5–7). For these aggregation assays, we incubated 1 mg/ml recombinant wild type TTR with increasing amounts of stabilizers at pH 4.3 and monitored protein aggregation both by absorbance at 400 nm and immuno-dot blot of the insoluble fraction, as we previously described (10). After 4 days in the absence of inhibitor, virtually the total amount of TTR present was converted to aggregates. In the presence of either tafamidis or diflunisal, TTR aggregation was diminished in a dose-dependent manner (Fig. 1a,b).

**Figure 1.**
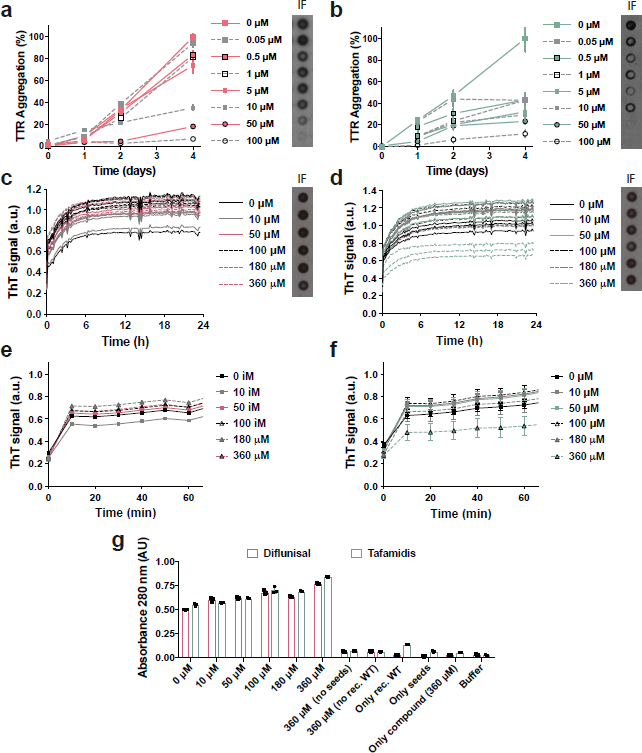
Tetramer stabilizers inhibit TTR aggregation in the absence of seeds and fail to inhibit amyloid seeding caused by ATTR-D38A *ex-vivo* seeds. (a, b) TTR aggregation in the absence of seeds measured by absorbance at 400 nm. Increasing amounts of diflunisal (a) or tafamidis (b) were added to 1 mg/ml recombinant wild type TTR and the sample was incubated for 4 days at pH 4.3. Absorbance measured after 4 days of incubation in the absence of ligand was considered 100% aggregation because no soluble TTR was detected. n=3, Error bars, SD. Right insets, anti-TTR dot blot of insoluble fractions (*IF*). (c, d) Amyloid seeding assay at pH 4.3, monitored by ThT fluorescence. Increasing amounts of diflunisal (c) or tafamidis (d) were added to 0.5 mg/ml recombinant wild type TTR and ATTR-D38A seeds. Insets, anti-TTR dot blot of insoluble fractions. All replicates are shown, n=3. a.u., arbitrary units. (e, f) Short-time view of lag phase of amyloid seeding assays in the presence of diflunisal (e) or tafamidis (f). n=3, Error bars, S.D. (g) 280 nm absorbance of insoluble fractions collected from *c* and *d*. AU, absorbance units.

We next studied the effect of various concentrations of tafamidis and diflunisal on protein aggregation in the presence of ATTR-D38A *ex-vivo* seeds. In our previous study, we observed that the addition of fibril seeds extracted from ATTR cardiac tissue accelerate aggregation of not only wild-type TTR at pH 4.3 but also monomeric TTR under physiological conditions (3). Additionally, we tested the effect of tafamidis and diflunisal at 180 µM on amyloid seeding and found that this concentration was not sufficient to hinder the process. Here we evaluate the effect of these ligands at various concentrations (Fig. 1c-f). For this assay, we incubated 0.5 mg/ml recombinant wild-type TTR with 30 ng/µl ATTRD38A *ex-vivo* seeds and increasing amounts of ligands. We monitored fibril formation for 24 hours by Thioflavin T fluorescence (ThT), by immuno-dot blot of the insoluble fraction (Fig. 1c-f), and by protein quantification of insoluble fractions (Fig. 1g), as we previously described (10). We found that these ligands did not reduce or delay seeded fibril formation even at ligand concentrations that resulted in full inhibition of TTR aggregation (Fig. 1).

### *Tetramer stabilization does not inhibit TTR amyloid formation induced by any of the ATTR* ex-vivo *seeds studied*

In our previous study, we report that our specimen obtained from an ATTR-D38A patient contains type B ATTR amyloid fibrils made of full-length TTR (3, 11). In order to rule out the possibility of pathology-based specificity, we analyzed seeding inhibition by ligands with seven additional ATTR samples (Fig 2). For this assay, we incubated 0.5 mg/ml recombinant wild-type TTR with 30 ng/µl ATTR *ex-vivo* seeds in the presence or the absence of 180 µM stabilizers. We monitored fibril formation by quantifying the protein content in the insoluble fraction after 24 hours of incubation at 37 ºC. Similarly, we found that the addition of tafamidis or diflunisal did not reduce the accumulation of insoluble material in the presence of seeds extracted from any of the other seven ATTR cardiac specimens. These findings suggest that tetramer stabilization by ligands is not an effective strategy to halt amyloid seeding under the studied conditions.

**Figure 2.**
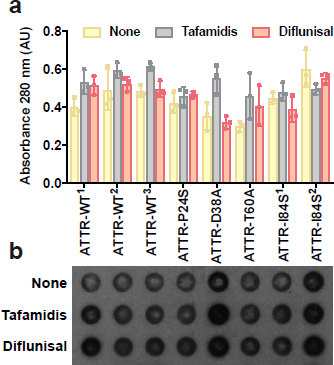
Tetramer stabilizers do not inhibit amyloid seeding caused by any of the cardiac samples analyzed. (a) Amyloid seeding assay of *ex-vivo* seeds incubated with recombinant wild type transthyretin in the absence or presence of 180 µM tafamidis or 180 µM diflunisal. After 24 h of incubation at pH 4.3, the insoluble fraction was collected, and absorbance was measured. AU, absorbance units. (b) Anti-TTR dot blot of insoluble fractions.

### T119M-derived stabilization fails to inhibit amyloid seeding

We then evaluated tetramer stabilization by mutagenesis, analyzing the effect of the T119M TTR variant on amyloid seeding. This variant exhibits high tetrameric stability that results in significant delay of the onset of familial neuropathic ATTR in patients who carry both *ttr*-V30M and *ttr*-T119M genes (5, 12, 13). Although a sample containing 1 mg/ml recombinant T119M variant remained soluble after days of incubation at a low pH that causes native TTR to dissociate (10) (Fig. 3a), the addition of 30 ng/µl ATTR-D38A *ex-vivo* seeds to 0.5 mg/ml recombinant T119M did result in seeded fibril formation (Fig. 3b). Notably, unlike T119M, the TTR variant T119W, which blocks self-association of strand H (10), did show a significant decrease of protein fibril formation upon seeding (Fig. 3b). We also found that although *ex-vivo* seeds caused the aggregation of a TTR variant that blocks self-association of the strand F, S85P/E92P, the lag phase was significantly longer (Fig. 3b-c). Consistent with our previous work (3), these findings suggest that blocking self-association of amyloidogenic TTR segments may be an effective approach to stop fibril formation when seeds are present.

**Figure 3.**
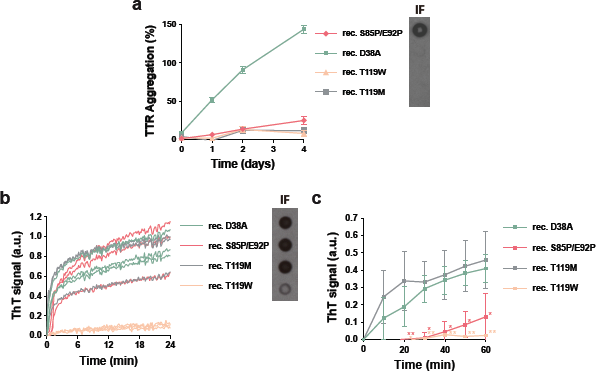
Mutational tetramer stabilization does not halt amyloid seeding. (a) Aggregation assay of 1 mg/ml non-aggregating recombinant TTR mutants in the absence of seeds at pH 4.3 followed by absorbance at 400 nm. Aggregation of wild type transthyretin was considered as 100%. Right inset, anti-TTR dot blot of insoluble fractions. n=3. Error bars, S.D. (b) Amyloid seeding assay of non-aggregating mutants in the presence of ATTR-D38A seeds, monitored by ThT fluorescence at pH 4.3. Right inset, anti-TTR dot blot of insoluble fractions. All replicates are shown, n=3. a.u. arbitrary units. (c) Short-time view of the lag phase of the amyloid seeding assay shown in b. n=3. Error bars, S.D. \**p* ≤ 0.05 and \*\**p* ≤ 0.005.

### *TabFH2 halts amyloid seeding caused by* ex-vivo *seeds extracted from all ATTR samples evaluated*

In a previous study, we developed peptide inhibitors that were designed to cap the tip of amyloidogenic segments of TTR in their amyloid state (10). Recently, we have shown that the optimized peptide inhibitor TabFH2 blocks amyloid seeding driven by fibrils extracted from a patient that carries the familial mutation ATTR-D38A (3). To better compare the efficacy of the peptide inhibitor TabFH2 with tafamidis and diflunisal, we evaluated TabFH2 with the same set of assays. We first found that TabFH2 inhibits TTR aggregation in the absence of ATTR seeds (Fig. 4a). For this assay, we incubated 1 mg/ml recombinant wild-type TTR with increasing amounts of TabFH2 and monitored protein aggregation for 4 days by absorbance at 400 nm, and visualized the insoluble fraction by immuno- dot blot, as described above. In addition, we analyzed the inhibitory effect of TabFH2 in our amyloid seeding assays, using the same conditions as for the previous experiment (Fig. 1c-f): we incubated 0.5 mg/ml of recombinant wild-type TTR with 30 ng/µl ATTR-D38A seeds and TabFH2 at various concentrations (Fig. 4b-d). We found that TabFH2 completely inhibits TTR amyloid seeding at concentrations higher than 180 µM with intermediate effect at lower doses. Altogether, our results indicate that TabFH2 inhibits TTR aggregation and amyloid seeding in a dose-dependent manner (Fig. 4).

**Figure 4.**
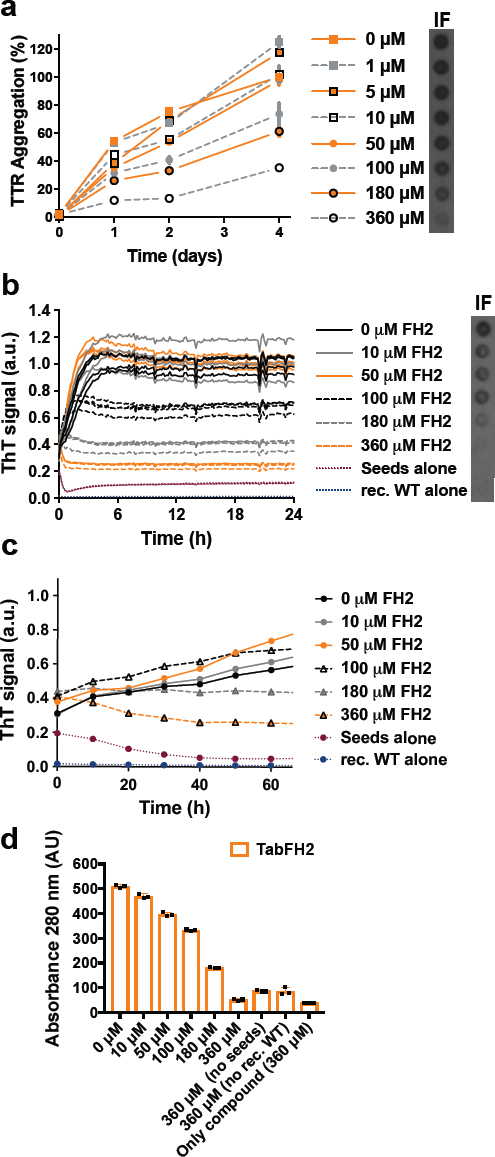
Tabfh2, An Anti-Amyloid Peptide-Based Inhibitor, Inhibits TTR aggregation and amyloid seeding caused by *ex-vivo* ATTR seeds extracted from ATTR-D38A cardiac tissue. (a) TTR aggregation in the absence of seeds measured by absorbance at 400 nm. Increasing amounts of TabFH2 were added to 1 mg/ml recombinant wild type TTR and the sample was incubated for 4 days at pH 4.3. Absorbance measured after 4 days of incubation in the absence of ligand was considered 100% aggregation because no soluble TTR was detected. n=3, Error bars, SD. Right insets, anti-TTR dot blot of insoluble fractions (*IF*). (b) Amyloid seeding assay at pH 4.3, monitored by ThT fluorescence. Increasing amounts of TabFH2 were added to 0.5 mg/ml recombinant wild type TTR and 30 ng/µl ATTR-D38A seeds. Insets, anti-TTR dot blot of insoluble fractions. All replicates are shown, n=4. a.u., arbitrary units. (c) Short-time view of lag phase of amyloid seeding assays in the presence of TabFH2. n=4, Error bars, S.D. (d) 280 nm absorbance of insoluble fractions collected from *b*. AU, absorbance units.

We next questioned the ATTR pathology specificity of TabFH2 by evaluating its inhibitory activity on our additional ATTR *ex-vivo* samples (Fig. 5). For this assay, we incubated 0.5 mg/ml recombinant wild-type TTR with 30 ng/µl ATTR *ex-vivo* seeds in the presence or the absence of 180 or 360 µM TabFH2. We monitored fibril formation by quantifying the protein content in the insoluble fraction after 24 hours of incubation at 37 ºC (Fig. 5a) and immuno-dot blot of the insoluble fraction (Fig. 5b). The presence or the absence of protein aggregates was confirmed by electron microscopy (Fig. 5c). A scrambled peptide, *H1*, was included as a negative control. We found that the efficacy of TabFH2 was dose-dependent, and although its efficiency differs from sample to sample, full inhibition was observed at the highest concentration analyzed (Fig. 5a-c). This variability did not show association with mutational background of the *ex-vivo* seeds. In contrast, we found an inverse relationship between TabFH2 effectiveness and the amount of TTR C-term fragments in the *ex-vivo* resuspension (Fig. 5d). The content of fragmented TTR, present in type A amyloidosis, was quantified from two independent western blots using an antibody that specifically recognizes TTR C-terminal fragments, as we previously described (3). Pearson’s correlation coefficient between amyloid seeding in the presence of 180 µM TabFH2 and TTR fragment content was of 0.90 (R-square=0.80) indicating a positive correlation.

**Figure 5.**
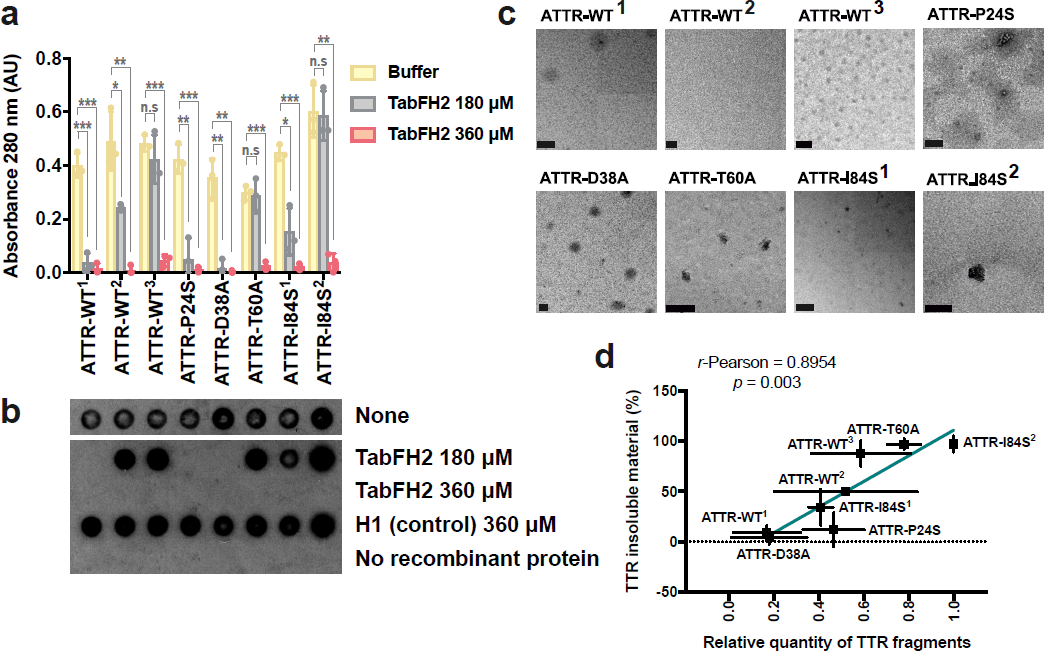
TabFH2 inhibits amyloid seeding caused by *ex-vivo* ATTR seeds extracted from all cardiac extracts. (a) Amyloid seeding inhibition evaluation. 30 ng/µl *ex-vivo* seeds were added to 0.5 mg/ml recombinant wild type transthyretin in the presence of 0, 180 or 360 µM TabFH2. Insoluble material was measured by absorbance at 280 nm of the insoluble fraction collected after 24 hours, and normalized to the absence of inhibitors to facilitate comparison of inhibitor performances. n=3. Error bars, S.D. \**p* ≤ 0.05 and \*\**p* ≤ 0.005, \*\*\**p* ≤ 0.0005. (b) TTR immuno-dot blot of insoluble fractions collected by centrifugation from the assay shown in a. (c) Electron micrograph of 0.5 mg/ml recombinant wild type TTR and 30 ng/µl *ex-vivo* seeds after 24 hours of incubation with 360 µM TabFH2. (d) Correlation between amyloid seeding capacity in the presence of 180 µM TabFH2, and relative quantity of truncated TTR of *ex-vivo* ATTR seeds. Truncated TTR content was quantified by ImageJ from two independent western-blots. Lineal regression and Pearson *r* were obtained by OriginLab.

### TabFH2 inhibitory activity is tissue independent

We extracted *ex-vivo* seeds from small quantities of two additional tissue types to evaluate tissue specificity of TabFH2. We extracted ATTR seeds from five samples of adipose tissue collected in fat pad biopsies, including one case of wild type ATTR and four cases of hereditary ATTR, and two samples of labial salivary glands from two ATTR-V30M patients (Table 1). Since biopsy specimens were considerably smaller than the cardiac tissue samples (30-150 mg vs 1-5 g), we downsized the amyloid extraction protocol accordingly. Otherwise, the procedure remained as previously described (3). The extraction from fat pad biopsies produced a very limited amount of insoluble material. We were therefore forced to reduce to 5 ng/µl the amount of *exvivo* seeds used in amyloid seeding assays. Due to the formation of structurally different species or perhaps a result of the small amount of seeds that were added to the assay, the ThT signal was insufficient to draw any conclusion. For this reason, we opted to follow amyloid seeding by image-based computational quantification of protein aggregates after 24 hours of incubation (Fig. 6). We collected images of the bottom of 96 well plates by optic microscopy using the Celigo S Imaging system. The images showed formation of UV-positive aggregates in those samples that contained *exvivo* seeds and no apparent aggregation in the control sample (Fig. 6a). The inhibitory effect of TabFH2 was confirmed by showing a significant reduction of amyloid conversion at 180 µM (Fig 6b). Together, our results indicate that TabFH2 efficiently inhibits amyloid seeding caused by wild type and mutant *ex-vivo* seeds in a tissue-independent manner.

**Figure 6.**
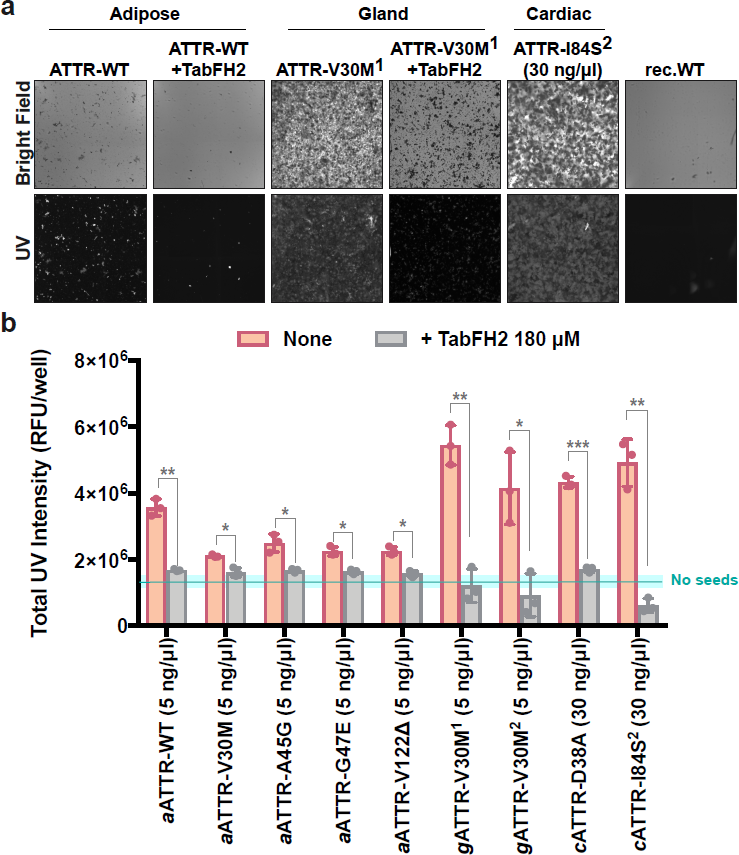
Inhibition of amyloid seeding caused by ex-vivo ATTR seeds extracted from fat pad and gland biopsies. (a) Representative optic micrographs of aggregates of recombinant wild-type TTR formed after 24 hours of incubation with 5 ng/µl fat-extracted ATTR-WT or gland-extracted ATTRV30M^1^ *ex-vivo* seeds, as visualized on a Celigo S Imaging system under bright field and UV channels. Two control samples are included: recombinant wild-type TTR aggregates formed after incubation with 30 ng/µl cardiac ATTR-D38A seeds, and recombinant wild-type TTR in the absence of seeds. (b) UV intensity-based quantification of protein aggregates in the presence and absence of 180 µM TabFH2. Blue line and rectangle represent the UV intensity mean and S.D. range in the absence of seeds, respectively. n=3. Error bars, S.D. \**p* ≤ 0.05 and \*\**p* ≤ 0.005, \*\*\**p* ≤ 0.0005, for the comparison between samples with and without TabFH2.

**Table 1.**
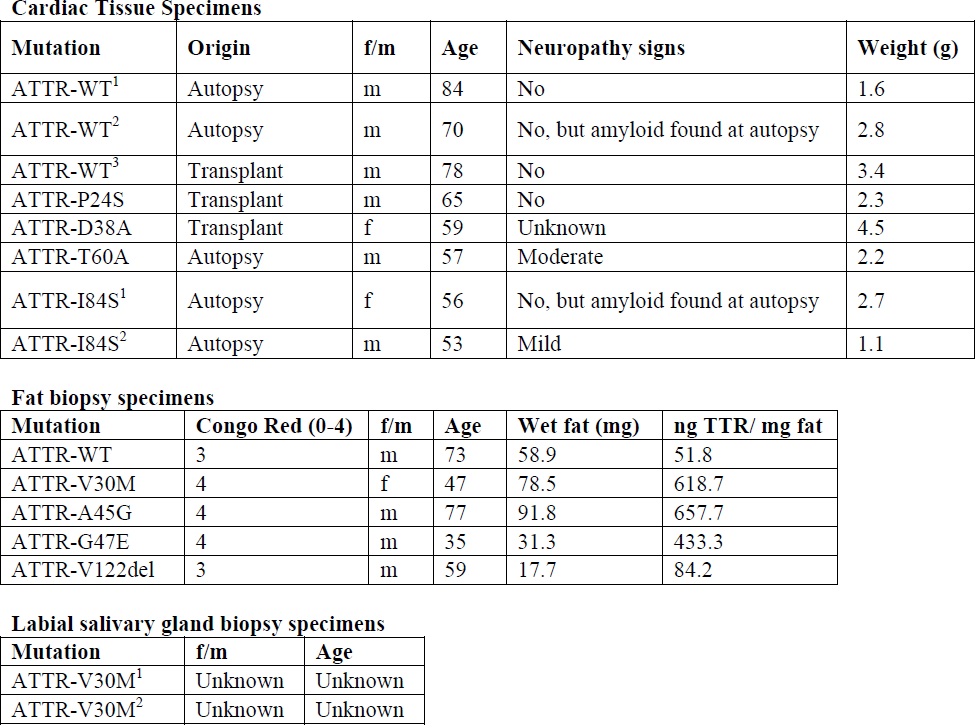
List of tissue specimens used for amyloid extraction. A total of 8 cardiac, 5 adipose, and 2 glandular specimens were included in this study. Various patients presented the same TTR variant, noted with superscript numbers.

Cardiac Tissue Specimens
| Mutation | Origin | f/m | Age | Neuropathy signs | Weight (g) |
| --- | --- | --- | --- | --- | --- |
| ATTR-WT<sup>1</sup> | Autopsy | m | 84 | No | 1.6 |
| ATTR-WT<sup>2</sup> | Autopsy | m | 70 | No, but amyloid found at autopsy | 2.8 |
| ATTR-WT<sup>3</sup> | Transplant | m | 78 | No | 3.4 |
| ATTR-P24S | Transplant | m | 65 | No | 2.3 |
| ATTR-D38A | Transplant | f | 59 | Unknown | 4.5 |
| ATTR-T60A | Autopsy | m | 57 | Moderate | 2.2 |
| ATTR-I84S<sup>1</sup> | Autopsy | f | 56 | No, but amyloid found at autopsy | 2.7 |
| ATTR-I84S<sup>2</sup> | Autopsy | m | 53 | Mild | 1.1 |

Fat biopsy specimens
| Mutation | Congo Red (0-4) | f/m | Age | Wet fat (mg) | ng TTR/ mg fat |
| --- | --- | --- | --- | --- | --- |
| ATTR-WT | 3 | m | 73 | 58.9 | 51.8 |
| ATTR-V30M | 4 | f | 47 | 78.5 | 618.7 |
| ATTR-A45G | 4 | m | 77 | 91.8 | 657.7 |
| ATTR-G47E | 4 | m | 35 | 31.3 | 433.3 |
| ATTR-V122del | 3 | m | 59 | 17.7 | 84.2 |

| Mutation | f/m | Age |
| --- | --- | --- |
| ATTR-V30M<sup>1</sup> | Unknown | Unknown |
| ATTR-V30M<sup>2</sup> | Unknown | Unknown |

### TabFH2 inhibits amyloid seeding by binding to ATTR fibrils

Our peptide inhibitors were originally designed and optimized to cap the tip of TTR fibrils by binding β-strands F and H (3, 10). To validate our design, we next assessed binding of TabFH2 to ATTR seeds (Fig. 7). We first immobilized 5 µg of ATTR-D38A seeds on each of eight anti-TTR pre-coated wells. Binding of ATTR seeds to the bottom of the wells was confirmed later by BCA protein assay of unbound material. We then added increasing amounts of TabFH2 (0-500 µM) to pretreated wells. After 2 hours of incubation, we analyzed the remaining amount of TabFH2 in the sample by HPLC. Anti-TTR pre-coated wells treated with buffer, but not seeds, were used as negative control to detect unspecific binding of TabFH2 to the well. While the majority of TabFH2 remained in the sample after incubation in negative control wells, most of TabFH2 was absent when wells were pretreated with ATTR seeds (Fig. 7d). Our results indicate that the mechanism of action of our peptide inhibitors involves binding ATTR fibrils.

**Figure 7.**
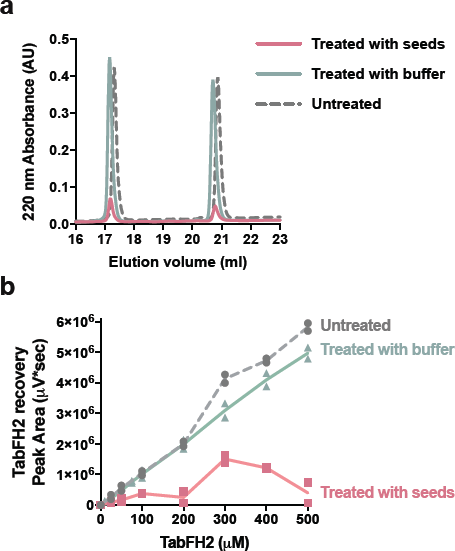
TabFH2 binds ATTR *ex-vivo* seeds. Increasing amounts of TabFH2 were added to independent wells pretreated with ATTR seeds (*treated with seeds*) or buffer (*treated with buffer*), and unbound TabFH2 was detected by reverse-phase chromatography. TabFH2 samples before treatment were included in the analysis (*untreated*). (A) Representative elution profile for a TabFH2 concentration of 500 µM. (B) Analysis of TabFH2 recovery at all concentrations.

## Discussion

Extensive studies by others have established that ligands such as tafamidis and diflunisal bind within TTR and stabilize its tetrameric structure, diminishing its rate of dissociation and hence its conversion to amyloid fibrils (5–7). The data of Fig. 1*a* and 1*b* confirm the stabilization of TTR by tafamidis and diflunisal. Indeed tafamidis has been prescribed for the treatment of neuropathic ATTR-V30M, and ameliorates disease progression when administered at disease stage I (14–17). Diflunisal has shown positive neurological effects in ATTR patients at different stages (9, 18). Despite the stabilization, our results show that disease-related seeds convert wild type TTR into amyloid fibrils in the presence of these ligands at concentrations that fully inhibit aggregation of recombinant transthyretin in the absence of seeds (Fig. 1 and 2) (19, 20). Thus our results offer a plausible explanation for the reported limited efficacy of tafamidis over the long term, when administered at late stages, or in patients with severe cardiac involvement (8, 21, 22). That is, stabilization of tetrameric TTR may be insufficient in situations in which seeded polymerization dominates rather than *de-novo* nucleation of TTR seeds.

Genetically stabilized TTR fails to inhibit seeded fibril formation. The genetic variant T119M of TTR and its capacity to delay fibril formation was originally found in a ATTR-V30M family because of its protective effects; this variant remains soluble at pH 4.4 for weeks, if not months (Fig. 3a) (10, 12, 23). However, in our experiments this stabilized variant does not halt conversion to amyloid in the presence of *ex-vivo* seeds (Fig. 3b, c). These findings may explain why heterozygous individuals carrying both the familial amyloidogenic *ttr*-V30M allele and the stabilizing *ttr*-T119M allele in time develop ATTR (12, 13). In ATTR-V30M/T119M patients, we hypothesize that the mutation T119M may delay the progression of ATTR by reducing *de-novo* nucleation.

The limited effect of TTR tetramer stabilization in our experiments does not contradict the well-established mechanism of *de-novo* formation of TTR amyloid (24). In our previous study, we show that the conversion of TTR to amyloid fibrils requires the dissociation of tetramers and partial unfolding of monomers by lowering the pH to 4.3, and only the monomeric variant MTTR can be seeded at physiological pH (3). In our assays, amyloid seeding is also performed at low pH to weaken the quaternary structure and to lead to tetramer dissociation. Under these conditions, the addition of seeds to recombinant TTR causes acceleration of fibril formation through a seeded polymerization. We reason that this seeded polymerization results in fibrils that are resilient to disassembly, thereby making the pathway irreversible. Because the interaction of TTR with the tetramer stabilizers is not irreversible, the presence of seeds will consequently and eventually reduce both monomeric and tetrameric pools while generating irreversible amyloid fibrils.

Capping of TTR amyloid fibrils by designed peptides is an effective approach to inhibit TTR fibril seeding. Our results show that in cases for which stabilization of tetrameric TTR by ligands may not be fully effective in halting fibril formation (Fig. 1 and 2), capping of TTR fibrils by designed blockers of fibril elongation is effective (Fig. 4-6). This may be of special importance for cardiac ATTR patients, who are often diagnosed when manifesting advanced TTR deposition and have limited treatment options. In the conditions of our experiments, TabFH2 blockers are effective in halting fibril formation caused by wild type ATTR seeds and also by seeds of at least five disease-related variants extracted from cardiac specimens (Fig. 5). Overall, we find that the inhibition of amyloid seeding by peptide inhibitors is an effective strategy independent of pathological variant or tissue (Fig. 5).

Structure-based inhibition of amyloid aggregation by peptides is an emergent strategy that has shown promising results. Our structure-based strategy has generated peptide inhibitors of tau fibril formation that is associated with Alzheimer’s disease (25, 26), SEVI amyloid aggregation that enhances HIV transmission (26), and p53 aggregation associated with certain types of ovarian cancer (27). In our previous work on inhibition of TTR, we generated TTR peptide inhibitors that block both protein aggregation and amyloid seeding catalyzed by *ex-vivo* ATTR fibrils (3, 10). Moreover, diseased flies showed motor improvement and a reduction of TTR deposition after treatment with our peptide inhibitors (28). In this study we expand the evaluation of peptide inhibitors to question their inhibitory capacity in comparison to two tetramer stabilizers and mutagenesis-based stabilization. Our results indicate that the inhibition of amyloid seeding by peptide inhibitors may represent a potential therapeutic strategy for ATTR when tetramer stabilization is not sufficient to halt disease progression.

## Experimental procedures

### Antibodies

The antibodies used in this study include rabbit anti-human transthyretin polyclonal antibody (DAKO, Agilent Technologies; immuno-dot blots and western blots 1:10,000) and horseradish peroxidaseconjugated goat anti-rabbit IgG antibody (ThermoFisher Scientific, immune-dot blots and western blots 1:5,000). Anti-truncated TTR was generously provided by Gunilla Westermark (labelled as 1898, western blots 1:5,000).

### Patients and Tissue Materials

Twenty ATTR patients carrying wild-type (n=4) or TTR mutations (n=16) were included in this study (refer to Table 1 for full details). Cardiac tissue specimens were obtained from several laboratories from explanted hearts or by autopsy. Adipose tissues were obtained from needle biopsy procedures performed at the University Medical Centre Groningen. Salivary gland tissues were obtained from surgical biopsy procedures performed at the Hospital Santo António in Porto. The UCLA Office of the Human Research Protection Program granted exemption from IRB review because all specimens were anonymized.

### Extraction of amyloid fibrils from tissue samples

Amyloid fibrils were extracted from fresh-frozen human tissue following a previously described protocol (3). In short, thawed tissue sample, resuspended in 10 mL of 0.15 M NaCl, was minced with a motorized homogenizer and pelleted by centrifugation at 15,000 rpm for 30 minutes. The pellet was subject to further cycles of resuspension, homogenization and centrifugation first with NaCl solution 7 times, followed by distilled water 3 times. Since there was less starting material for both adipose tissue and salivary gland specimens, the extraction protocol was downsized in volume accordingly. The final pellet was lyophilized and amyloid content of the extracts was confirmed by electron microscopy. TTR content of the samples was analyzed by anti-TTR western blot.

### Purification of Recombinant TTR

Recombinant transthyretin was prepared as described previously (10). To summarize, E. coli (Millipore Rosetta DE3 pLysS Competent Cells) were transformed with a pET24(+) vector carrying the sequence for either wild-type or a mutant of transthyretin. The expressed recombinant protein was harvested and purified by nickel affinity chromatography with a HisTrap column (GE Healthcare). The appropriate fractions were pooled and further purified by size exclusion on a Superdex 75 gel filtration column (GE Healthcare). Recombinant transthyretin was stored in 10 mM sodium phosphate pH 7.5, 100 mM KCl, and 1 mM EDTA at −20 °C.

### Western Blot of Tissue Extracts

TTR content of tissue extracts was confirmed by western blotting as described previously (3). To summarize, equal amounts of total protein were loaded on a 4-12% NuPAGE BisTris Gel (ThermoFisher Scientific) and separated by gel electrophoresis in denaturing conditions. TTR was detected after transfer to a nitrocellulose membrane with polyclonal anti-human transthyretin antibody or anti-truncated TTR antibody and horseradish peroxidase conjugated secondary goat anti-rabbit IgG. SuperSignal^TM^ West Pico Chemiluminescent Substrate (ThermoFisher Scientific) was used according to manufacturer’s instructions to visualize TTR content. Truncated TTR content of cardiac seeds were quantified by ImageJ using two independent western blots.

### Congo red staining and TTR content quantification of adipose tissue

Abdominal fat smears were made as previously described (29). Slides were stained with alkaline Congo red (30) and apple-green birefringence under polarized light were semi-quantitatively scored as follows: 0 (negative), 1 (minute, <1% surface area), 2 (little, between 1 and 10%), 3 (moderate, between 10 and 60%), and 4 (abundant, >60%). The remaining abdominal fat tissue was weighted and washed. Proteins were resuspended with a Tris- guanidinium hydrochloride solution, and TTR content was measured by ELISA. Briefly, microtiter plates were coated overnight with the extracts and human native TTR protein (Abcam, Cambridge, UK), which served as control, in several dilutions. Detection was done by using rabbit anti-human TTR polyclonal antibodies (DAKO, Agilent Technologies) followed by horseradish peroxidase- conjugated goat anti-rabbit IgG antibody (DAKO, Agilent Technologies) and visualized by a color reaction with TMB ELISA substrate. Plates were scanned at 450 nm after stopping the reaction with sulfuric acid.

### Non-seeded TTR Aggregation

TTR aggregation assays were done as previously described (10). Briefly, 1 mg/mL TTR sample in 100 mM sodium acetate pH 4.3, 100 mM KCl, 1 mM EDTA was incubated in the presence or absence of inhibitor – diflunisal, tafamidis or TabFH2 – at 37°C for 4 days. TTR aggregation was followed by measuring sample turbidity at 400 nm and by anti-TTR immuno-dot blot of the insoluble fraction.

### TTR Amyloid Seeding Assay

Extracted tissue samples were used to seed the formation of transthyretin amyloid fibrils following a protocol described previously (3). In short, extracts were washed twice in 1% sodium dodecyl sulfate and twice in 10 mM sodium phosphate pH 7.5, 100 mM KCl, 1 mM EDTA. Next, the extracts were sonicated at minimum intensity with 5 second pulses for a total of 10 minutes. Protein concentration of the samples was determined by Pierce^TM^ BCA Protein Assay Kit (ThermoFisher Scientific). Seeding reactions contained 0.5 mg/mL of recombinant protein, 30 ng/µL of cardiac extract or in the case of adipose and gland extracts 5 ng/µL, 5 µM thioflavin T, 100 mM sodium acetate pH 4.3, 100 mM KCl, and 1 mM EDTA. The inhibitors mentioned, diflunisal, tafamidis, and TabFH2 were added at concentrations described in the figures. Thioflavin T fluorescence emission was measured at 482 nm with excitation at 440 nm in a FLUOstar Omega (BMG LabTech) plate reader. Plates were incubated at 37°C for 24 hours with orbital shaking at 700 rpm between measurement points. In all assays, measurements were normalized by subtracting the initial ThT measurement and considering the maximum signal as 100%. Protein aggregates were visualized by both bright field and UV and TTR aggregation was quantified using a Celigo S Imaging system. The insoluble fraction was obtained by centrifuging the sample and resuspending the pellet in guanidinium hydrochloride. Protein content of the pellet was determined by measuring absorbance at 280 nm.

### Anti-TTR immuno-dot blot

TTR aggregation was confirmed by immuno-dot blot as mentioned previously (10). After obtaining the insoluble fraction by centrifugation and resuspension in guanidinium hydrochloride, 15 µL of sample was dotted onto a nitrocellulose membrane (0.2 µm, Bio-Rad). TTR content was visualized using polyclonal rabbit anti-human transthyretin (DAKO), horseradish peroxidase conjugated goat anti-rabbit IgG antibody (ThermoFisher Scientific), and SuperSignal^TM^ West Pico Chemiluminescent Substrate (ThermoFisher Scientific).

### Transmission Electron Microscopy

Amyloid content of tissue extracts and inhibition of fibril formation was confirmed by transmission electron microscopy. 5 µL of sample was applied to a glow discharged carbon coated EM grid (CF150-Cu, Electron Microscopy Sciences) for 4 minutes. After three quick rinses in distilled water, grids were stained with 2% uranyl acetate for 2 minutes. Samples were visualized using a T12 Quick CryoEM and CryoET (FEI) transmission electron microscope using an acceleration voltage of 120 kV equipped with a Gatan 2,048 x 2,048 CCD camera.

### Detection of TabFH2 binding to ATTR seeds

TabFH2 binding to ATTR seeds was analyzed by HPLC. First, ATTR-D38A fibrils were immobilized on anti-TTR pre-coated well plates (Prealbumin ELISA kit, Abcam) as follows. 50 ml samples containing 0.1 mg/ml ATTR seeds were added on each well. Control wells were equally treated with buffer (10 mM sodium acetate pH 7.5, 100 mM KCl, 1 mM EDTA). After 1 hour incubation at room temperature, samples were then removed and wells were washed twice with buffer. Immobilization of the total amount of fibrils was confirmed by BCA protein assay of the remaining sample. 50 ml samples that contained increasing concentrations of TabFH2 (0-500 mM) were added to independent wells pretreated with ATTR seeds or buffer. After an incubation of 2 hours at room temperature, samples were transferred to new tubes and snap frozen until further analysis. Unbound TabFH2 was detected by chromatography after 0.10 nm filtration, on an Waters 1525 HPLC System (SpectraLab), with a Proto 300 C18 5 mm 250 x 4.6 mm analytical reverse phase column (Higgins Analytical). Flow rate = 1.0 ml/min; solvents: A = 0.1% trifluoroacetic acid in water, and B = 0.1% trifluoroacetic acid in acetonitrile. The column was equilibrated with 10% B for 5 minutes, followed by a gradient from 10% to 60% B in 30 minutes, and a 2 minute wash at 95% B. TabFH2 eluted in two peaks after approximately 17 and 20 min from start. Peaks were integrated by Breeze2 software and Prism was used for graphing.

## Acknowledgments

We thank Dr. Jeffery Kelly for a gift of tafamidis and discussions, Drs. Joel Buxbaum and Duilio Cascio for helpful discussion, the Amyloidosis Foundation (Grant #20170827), National Institutes of Health (Grant #R01- AG048120), and The Howard Hughes Medical Institute for support, and to the patients who generously donated tissues.

## Conflict of interest statement

The authors and UCLA have filed an international patent application for the TTR inhibitors (No. PCT/US17/40103). D.S.E. is an advisor and equity holder of ADRx, Inc. L.S. is a consultant of ADRx, Inc.

